# Minimal impact of ivermectin on immune response and transcriptional profiles in naïve adults with mild COVID-19

**DOI:** 10.1101/2025.03.31.646276

**Authors:** Marta Ribes, Cèlia Torres, Mar Canyelles, Rocío Rubio, Marta Vidal, Luis Izquierdo, Andrés Blanco-Di Matteo, Iñigo Pineda, Alejandro Fernandez-Montero, Carlota Jordan-Iborra, Francisco Carmona-Torre, José R Yuste, Jose L Del Pozo, Gabriel Reina, Belen Sadaba, Mirian Fernández-Alonso, Pere Santamaria, Carlo Carolis, Ruth Aguilar, Dídac Macià, Carlos Chaccour, Carlota Dobaño, Gemma Moncunill

## Abstract

Ivermectin (IVM), an antiparasitic drug, was repurposed to treat COVID-19 based on its in vitro antiviral effects. However, it was abandoned after multiple clinical trials reported a lack of efficacy. Immunomodulatory effects have been proposed but remain unclear, yet they may be relevant given IVM use for other infections. We assessed the IVM immunomodulatory effect in 24 participants from a clinical trial evaluating its potential to reduce COVID-19 transmission in mild cases within 48 hours of symptoms onset. The IVM-treated patients showed non-significant lower viral loads, and a significantly shorter duration of hyposmia/anosmia. We measured IgG, IgA, and IgM against five SARS-CoV-2 antigens, and 30 cytokines by Luminex, alongside whole blood RNA sequencing, pan-leukocyte immunophenotyping, and SARS-CoV-2-specific T cell analysis by flow cytometry from day 1 to day 28 post-treatment. All antibody responses increased from day 4, while 13 cytokines significantly decreased over time (adjusted p<0.05). IVM-treated patients had only significantly higher anti-nucleocapsid IgG levels at day 4 (adjusted p=0.041) and 7 (adjusted p=0.045) compared to placebo. SARS-CoV-2-specific CD4^+^ and CD8^+^ T cells increased over time, with significantly higher effector memory CD4^+^ T cells at day 7 compared to day 1 (p=0.027) and the only difference between groups was lower frequencies of spike-specific naïve CD4^+^ T cells at day 7 in IVM-treated participants (0.006% vs 0.036% p=0.02). Transcriptomic data showed downregulation of innate and antiviral blood transcriptional modules (BTMs) over time, with an increase in adaptive immune related BTMs. While no differential gene expression was detected, the IVM-treated had upregulated innate and downregulated T cell and cell cycle BTMs compared to placebo. Overall, our comprehensive longitudinal analysis of early immune responses in mild COVID-19 revealed no robust immunological effects of IVM, consistent with clinical trials results and suggesting a lack of efficacy of IVM in COVID-19 treatment.

**AUTHOR SUMMARY:** Ivermectin (IVM), an antiparasitic drug, was tested in clinical trials as a potential treatment for COVID-19 due to its in vitro antiviral properties and hypothetical immunomodulatory effects. In this study, we explored the immunomodulatory effects of IVM in a clinical trial involving 24 mild COVID-19 patients who received IVM within 48 hours of symptoms onset. The IVM group showed a trend towards lower viral loads post-treatment, and a significantly shorter duration of hyposmia/anosmia. We comprehensively analyzed immune responses by measuring antibody levels, cytokine profiles, immune cell subsets and whole blood gene expression over 28 days. The IVM group only had increased levels of IgG against the SARS-CoV-2 nucleocapsid compared to the control group. IVM did not affect the kinetics of SARS-CoV-2 T cells, despite a slight decrease in naive CD4^+^ T cells. Additionally, gene expression analysis showed a decrease in innate and antiviral responses and an increase of adaptive responses over time that were slightly stronger in the placebo compared to the IVM group. In conclusion, despite some differences in IVM treated participants, our detailed analyses do not support significant immunomodulatory effects that could benefit disease progression.

## INTRODUCTION

Ivermectin (IVM) is widely used as an endectoparasitic drug in human and veterinary medicine. However, it has also shown antiviral properties [1] and was tested as such at the beginning of the COVID-19 pandemic for potential use against SARS-CoV-2 infection. An early study reported an in vitro inhibition of SARS-CoV-2 proliferation but with a concentration 35-fold higher than that approved for clinical dosages in humans [2]. This finding led to numerous randomized clinical trials testing its efficacy. A Cochrane systematic review including 11 trials concluded that, for outpatients, there was low-to high-certainty evidence that IVM had no beneficial effect for people with mild-to-moderate COVID-19 [3]. For inpatients, there was low-certainty evidence that IVM offered no benefit in terms of disease progression, prevention of death, viral clearance and adverse events [3].

The mode of action of IVM is not fully understood, nor is the nature of its wide range of effects in many different organisms beyond its endectoparasitic properties [4]. It has been long thought that the activity of IVM may not be restricted to the neuro-physiology of parasites (through its selective binding to glutamate-gated chloride ion channels), and that it may also modulate the immune system of animals and humans [5]. This immunomodulation could be indirect, through the inhibition of the parasite mechanisms of immune evasion, or direct, through changes in immune function independently of infection. However, literature on the topic is inconsistent [5].

In parasite infected lambs, IVM treatment led to less blastogenic activity and antibody levels [6]. In mites infested rabbits, IVM increased antibody production, but it was interpreted as a response to the massive and sudden death of mites [7]. CD8^+^ T cells decreased with time in Capra hircus papillomavirus (ChPV)-infected goats treated with IVM, but not in the control group [8]. In humans, IVM studies were mainly carried out in onchocerciasis and filariasis patients, finding a potentiation of the cellular component [9,10] and cytokine concentrations [9,11] as well as a decrease in antibody levels [10,11]. Among COVID-19 convalescent patients taking self-prescribed IVM, a dose-dependent decrease in antibody production by peripheral blood mononuclear cells (PBMCs) was found when stimulated with SARS-CoV-2 spike (S) protein, with an increase in reported symptoms, while the T-helper (Th)1/Th2 balance was not affected by IVM dosage [12].

In uninfected laboratory animals, an increase in antibody levels was found in mice and rabbits when treated with IVM and exposed to exogenous antigens [13,14]. In addition, an IVM dose-dependent increase in macrophage engulfment of sheep red blood cells in rabbits was observed [15]. Topical IVM administration decreased T-cells and cytokines in a murine skin allergy model [16]. In vitro, IVM decreased the inflammatory cytokine production by lipopolysaccharide (LPS)-stimulated macrophages [17] and an inhibitory effect of IVM was also observed on T cell proliferation and functions [16]. In humans, a 0.25 mg/kg single IVM dose in healthy volunteers did not significantly change cytokine levels and immune cell gene expression compared to placebo recipients [18].

In addition to its immunomodulatory effects, IVM has been explored for its potential beneficial impact on clearing SARS-CoV-2 infection via alternative mechanisms, as discussed in several reviews [19,20]. IVM disrupts the association of importin (IMP)α and IMPβ, impeding the nuclear import required for replication of viruses like HIV-1 and Dengue [21]. It has been suggested that this impairment of nuclear transport of viral particles might be beneficial against SARS-CoV-2 pathogenesis, hindering its immune response evasion and replication [19,20]. Some studies have suggested that IVM might disrupt SARS-CoV-2 binding to angiotensin-converting enzyme 2 (ACE2) receptor [22,23], which would hamper the host cell infection. Another suggested mode of action comes from IVM high affinity for RNA-dependent RNA polymerases (RdRp) [24], an essential enzyme for viral replication. However, the affinity or inhibition has not been tested for the SARS-CoV-2 RdRp specifically.

Therefore, despite the IVM antiviral activity at higher concentrations of those recommended for human clinical use, the potential immunomodulatory properties of IVM could still benefit COVID-19 disease progression. Here, we aimed to assess the effect of IVM on the innate and adaptive immune responses to SARS-CoV-2 in early symptomatic infections. We performed an ancillary study to the SARS-CoV-2 Ivermectin Navarra-ISGlobal Trial (SAINT)[25,26], a clinical trial designed to evaluate the potential of IVM (single dose of 400 mcg/kg) to reduce COVID-19 transmission in low-risk non-severe patients within the first 48 h of symptoms onset. The trial did not find a significant difference for the primary endpoint between treatment arms, but the IVM group presented a non-significant trend towards lower viral loads at days 4 and 7, and had significantly less days of reported hyposmia/anosmia (76 vs 158-patient-days) [26].

## RESULTS

### Baseline characteristics of participants

A total of 24 patients with mild COVID-19 participated in the SAINT double-blind, placebo-controlled, randomized clinical trial [26]. Twelve received IVM and twelve, placebo. The two treatment arms were balanced with regards to age (median age 26 in both groups), sex (42% females in IVM group and 58% in placebo group), baseline viral load (3.7 x 10^8^ N gene copies/mL in IVM group and 3.3 x 10^8^ in placebo group), hours since symptoms onset (median 24 h (IQR 24-48) in IVM-group and 48 h (IQR 36-48) in placebo group), body mass index (median 23.5 in IVM group and 22.9 in placebo group) and classical lab measurements, except for the mean systolic (113.5 vs 125.8, F=11.6) and diastolic (75.2 vs 81.6, F=6.0) pressures, which were significantly higher in the placebo-treated group. Values for baseline measurements were presented in the paper reporting the clinical trial results [26].

Exclusion criteria to participate in the clinical trial mandated that at day of drug intake (day 1), participants had non-detectable IgG levels against SARS-CoV-2 by a rapid test, and no more than 72 h of fever or cough. None of the participants progressed to severe disease [26].

### Ivermectin effect on antibody responses to SARS-CoV-2

Levels of IgA, IgG and IgM to the nucleocapsid (N) and S SARS-CoV-2 antigens generally increased from day 4 post-treatment (Figure 1). Noteworthy, day 1 did not equal the first day of disease, it was on average 43 h since first symptom, which could be 4.5 days since infection according to the mean incubation period reported for the Wuhan strain [27]. A participant did not develop IgA responses. IgA and IgM peaked approximately at day 14 and waned thereafter, while IgG against N and S2 peaked abruptly at day 14 post-treatment and then plateaued. IgG levels to S, S1 and RBD did not seem to reach the peak within the 28-days follow-up duration, which is in line with the literature [28]. Levels of all immunoglobulin isotypes to other human coronaviruses (HuCoVs) N antigens remained virtually unchanged, indicating no cross-reactive responses (Supplementary Figure 1).

**Figure 1.**
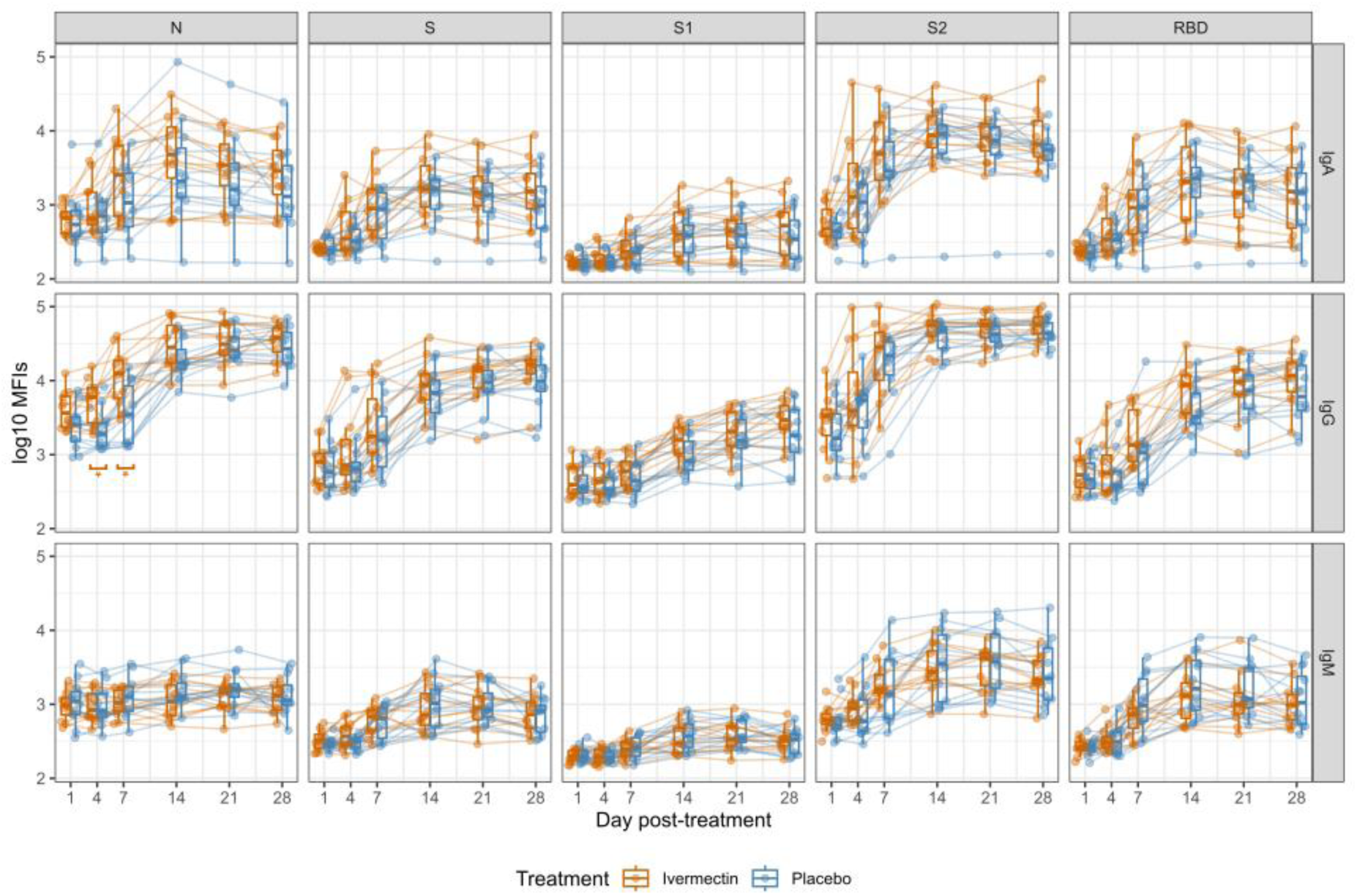
Kinetics of SARS-CoV-2-specific antibody levels since day of treatment. Log-10 transformed levels (median fluorescence intensity, MFI) of IgA, IgG, and IgM against each antigen (nucleocapsid full-length protein (N), the receptor binding domain (RBD), full length spike protein (S) and its subregions (S1 and S2) measured in 24 patients across six time points (data from same individual are joined by lines). For each time point boxplots represent median and interquartile range MFI for ivermectin-treated patients (orange) and placebo-treated patients (blue). The orange horizontal bars indicate significant differences in levels between treatment arms at a given time point. * Indicates adjusted p < 0.05 from Wilcoxon-test with Benjamini-Hochberg correction. Statistical differences in levels between timepoints are not shown.

At baseline or day 1 (treatment day), there were no significant differences between treatment arms for any of the antibody-antigen pairs. Only IgG levels to N exhibited significant differences in geometric means between IVM and placebo arms in subsequent time points. IVM-treated patients had significantly higher anti-N IgG levels at day 4 (mean log_10_-transformed mean fluorescence intensity (MFI) 3.71 vs 3.33, adjusted p = 0.041) and day 7 (4.04 vs 3.55, adjusted p = 0.045) than placebo participants. The rest of antigen-isotype pairs assessed did not show significant differences. However, there was a trend towards IVM-treated patients having higher IgG and IgA levels and lower IgM levels compared to placebo-treated patients, although this was already seen at baseline.

### Ivermectin effect on cytokine profiles

We analyzed a panel of 30 cytokines in plasma samples covering different immunological functions. We found a vast heterogeneity in their kinetics (Figure 2, Supplementary Figure 2). Thirteen of them decreased significantly over time (adjusted p<0.05) (Figure 2), with interferon (IFN)-γ induced protein (IP-10), interleukin (IL)-8 and IL-12 decreasing more steeply from day 1 and stabilizing at these lower levels from day 14 post-treatment. On the other hand, the pro-inflammatory cytokines IFN-α, IL-1β, IL-6, IL-7, and IL-15, the anti-inflammatory IL1-RA, and the chemokines eotaxin, monocyte chemoattractant protein (MCP-1) and monokine induced by IFN-γ (MIG), along with vascular endothelial growth factor (VEGF), exhibited a much smoother decay (Figure 2). This waning in cytokines follows the suppression of viral load at day 4 (Figure 3).

**Figure 2.**
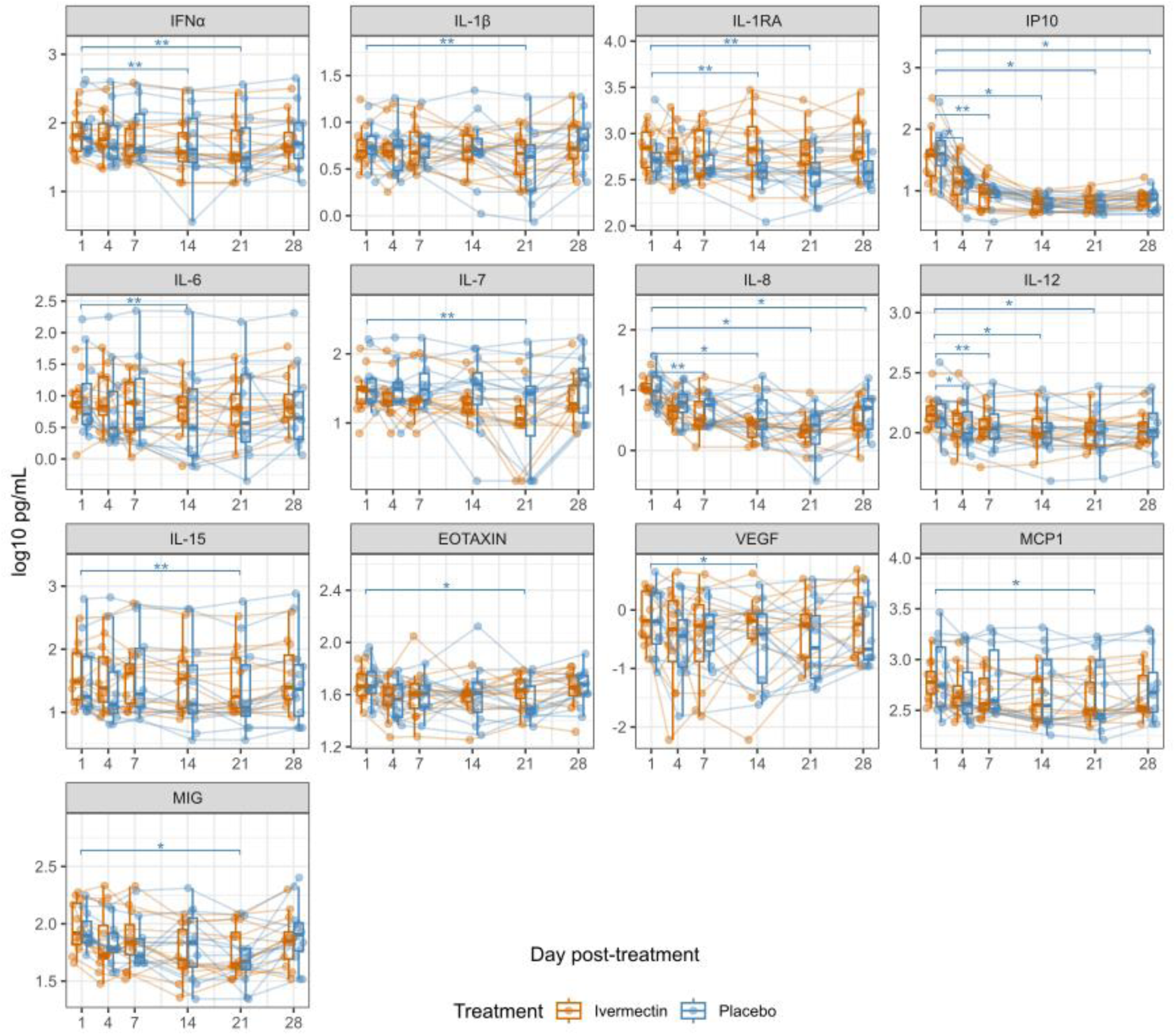
Kinetics of plasma cytokine concentrations since day of treatment. Log-10 transformed levels (pg/mL) of cytokines measured in 24 patients across six time points (data from same individual are joined by lines). For each time point boxplots represent median and inter quartile range pg/mL for ivermectin-treated patients (orange) and placebo-treated patients (blue). Here are represented the 13 out of the 30 cytokines analyzed that significantly varied over time after adjusting for multiple comparisons. The rest of cytokines are presented in Supplementary Figure 2. The blue horizontal bars indicate significant differences in levels between time points with regards to day 1 among placebo-treated patients (n=12). * Indicates adjusted p < 0.05, and ** adjusted p <0.01 from T-test with Benjamini-Hochberg correction.

**Figure 3.**
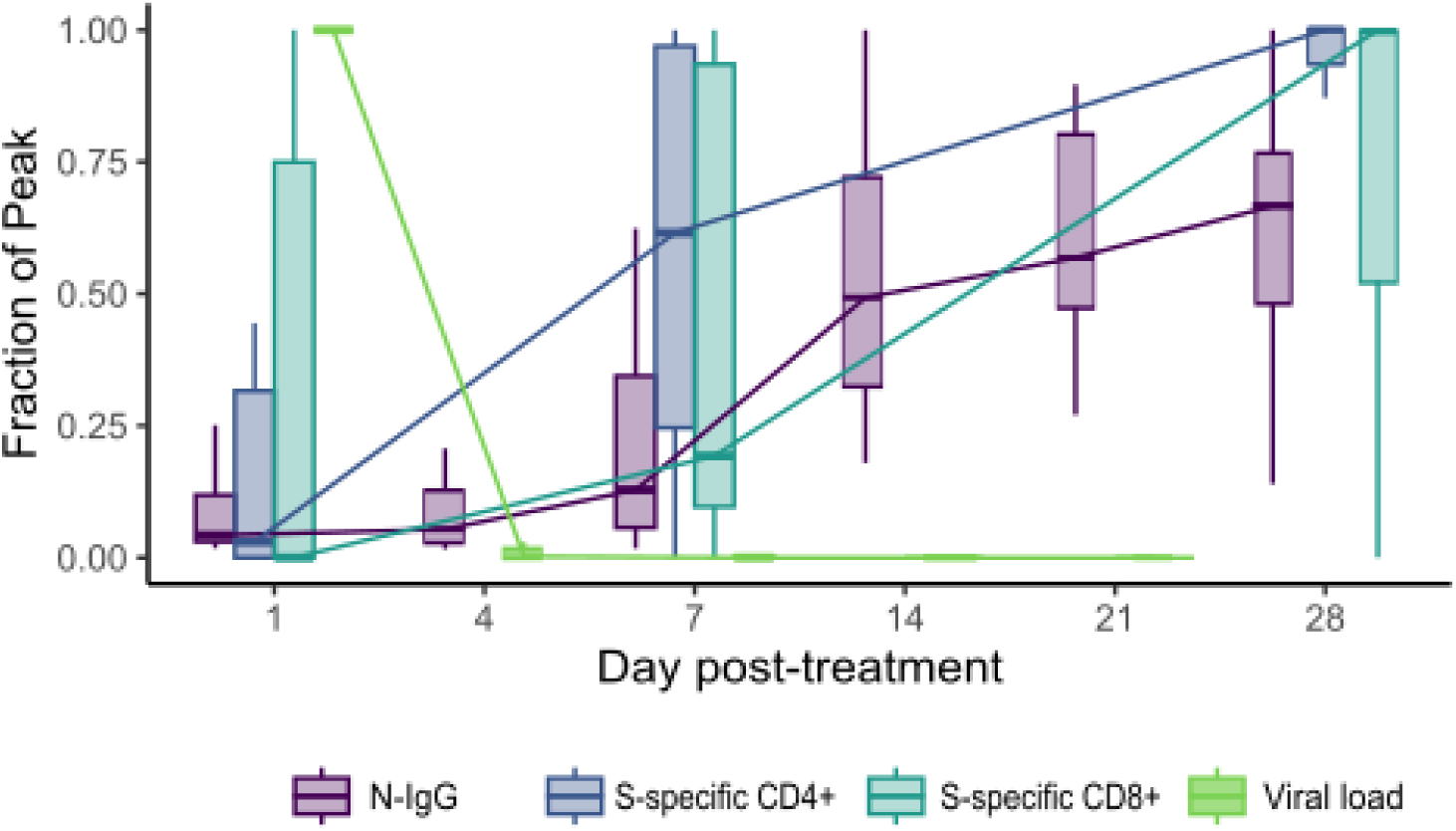
Overlaid kinetics of representative markers of the different immune response components described in other Figures. Levels across time points are depicted as the fraction of the peak response observed. N-IgG: IgG levels against the nucleocapsid (N); S-specific CD4^+^ are frequencies of spike specific CD4^+^ T cells within total CD4^+^ T cells; S-specific CD8^+^ are frequencies of spike specific CD8^+^ T cells within total CD8^+^ T cells (n=24).

Among all cytokines, only IL-1RA, an antagonist of the receptor of the proinflammatory cytokine IL-1β that is induced in response to IL-1β increases, had statistically significantly different levels between the two treatment arms. The IVM-treated group had higher levels of IL-1RA at day 4 (mean log_10_ (pg/mL) 2.8 vs 2.64, p=0.0462, p-adjusted=0.092), day 14 (2.89 vs 2.61, p=0.028, p-adjusted=0.092), day 21 (2.82 vs 2.59, p=0.032, p-adjusted=0.092) and day 28 (2.88 vs 2.65, p=0.028, p-adjusted=0.092), although these were not significant when adjusted for multiple comparisons.

### Ivermectin effect on SARS-CoV-2-specific T cells

To assess the cellular response to SARS-CoV-2 along the course of the infection, antigen-specific T-cells to the S, membrane (M) and N viral proteins were measured through the activation-induced markers (AIM) flow cytometry assay (Supplementary Figure 3). Activated antigen-specific CD4^+^ and CD8^+^ T cells were defined through the upregulation of the activation markers CD137 and OX40 or CD69, respectively, and their Th1 and Th2 profiles were defined through the expression of CXCR3 and CCR4, respectively. Antigen-specific cells were also assessed within each memory subset, defined by the expression of CCR7 and CD45RA.

As expected, S and MN-specific CD4^+^ T cells, and their memory phenotypes, increased progressively with time after infection (Figure 4, Supplementary Figure 4). S- and MN-specific CD8^+^ T cell increases where less pronounced and affected mainly effector memory re-expressing CD45RA (EMRA) and central memory (CM) subsets. There were no major changes in the frequency of antigen-specific cells expressing Th1/Th2 markers, which stabilized after day 7 (Figure 4 and Supplementary Figure 4). Statistically, these increases in mean frequencies were not significant after adjustment for multiple comparisons, except for S-specific effector memory (EM) CD4^+^ T cells, which were significantly higher at day 7 post-treatment with regards to day 1 (adjusted p=0.027).

**Figure 4.**
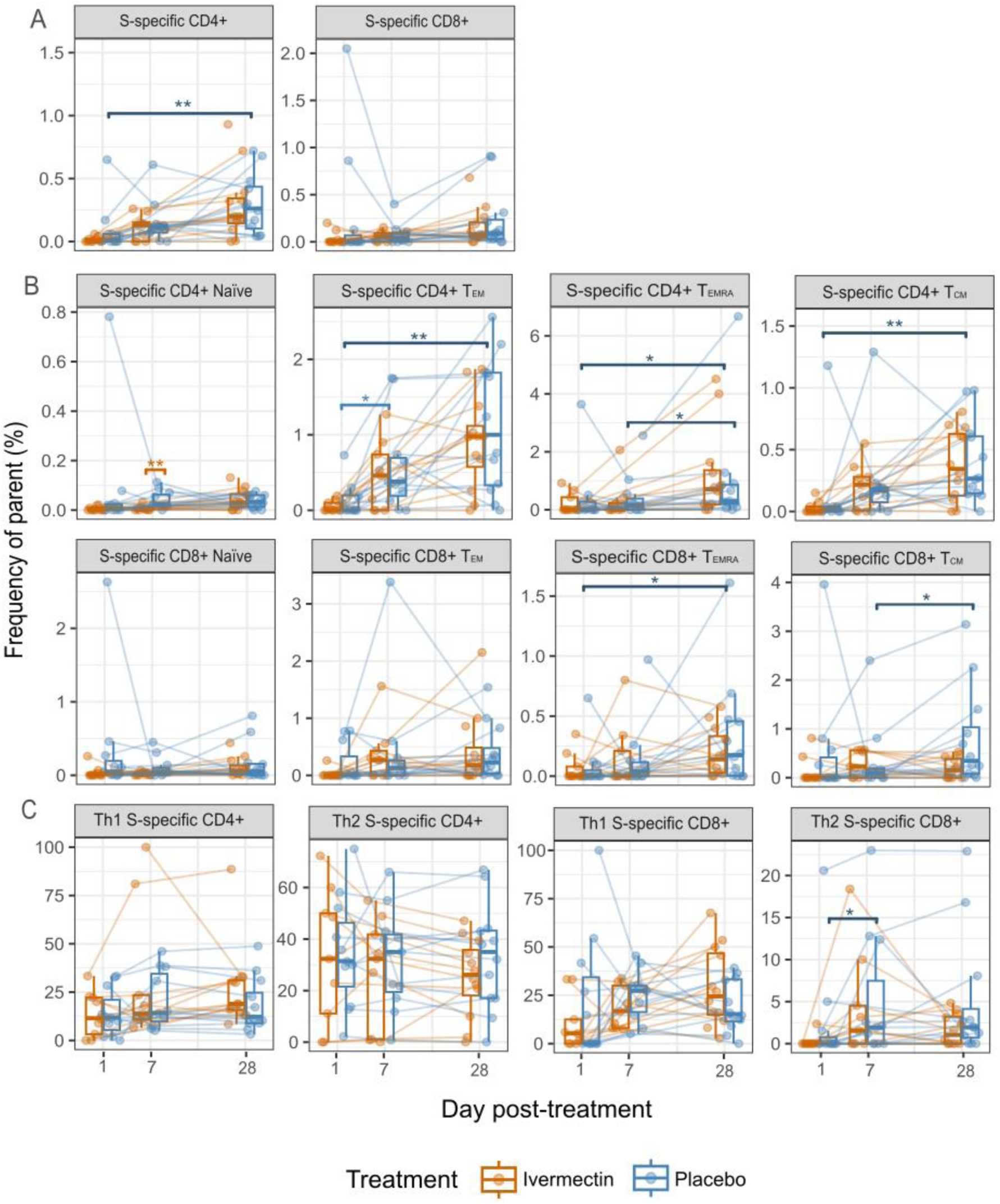
Kinetics of SARS-CoV-2-specific T cell populations since day of treatment. Frequencies of AIM^+^ CD4^+^ and CD8^+^ T cell subsets after spike stimulation and background subtraction. **A**. Frequencies (%) of S-specific CD4^+^ and CD8^+^ T cell populations. **B**. Frequencies (%) of S-specific CD4^+^ and CD8^+^ T cell memory subsets. **C.** Frequencies (%) of Th1 and Th2 within S-specific CD4^+^ and CD8^+^ T cells. For each time point box plots represent median and inter quartile range for 12 ivermectin-treated patients (orange) and 12 placebo-treated patients (blue), paired samples joined by lines. For day 1, a placebo-treated sample and three ivermectin-treated samples were not included in the assay. For day 7, another placebo-treated sample and three other ivermectin-treated samples were not included. Statistically significant differences between time points among placebo-treated patients before adjusting appear in dark blue brackets and * indicate p < 0.05, and ** p <0.01 from Wilcoxon-test (n=12). The lighter blue bracket indicates statistically significant difference after adjusting for Benjamini-Hochberg correction (*adjusted p <0.05) (n=12). Statistically significant differences between treatment arms before adjusting appear in orange brackets and ** indicates p <0.01 from Wilcoxon-test.

SARS-CoV-2 specific CD4^+^ T cells had predominantly a CM and EM phenotype (Figure 5), whereas for CD8^+^ T cells, the predominant phenotypes detected were EM and EMRA, in line with what has been reported in the literature [29]. No differences in phenotype were observed between S and MN specific T cells. Among specific CD4^+^ T cells, the Th2 profile predominated over Th1 (mean 58% of Th2 vs 42% of Th1 among S^+^ CD4^+^ T cells and 52.3% of Th2 vs 47.7% of Th1 among NM^+^ CD4^+^ T cells).

**Figure 5.**
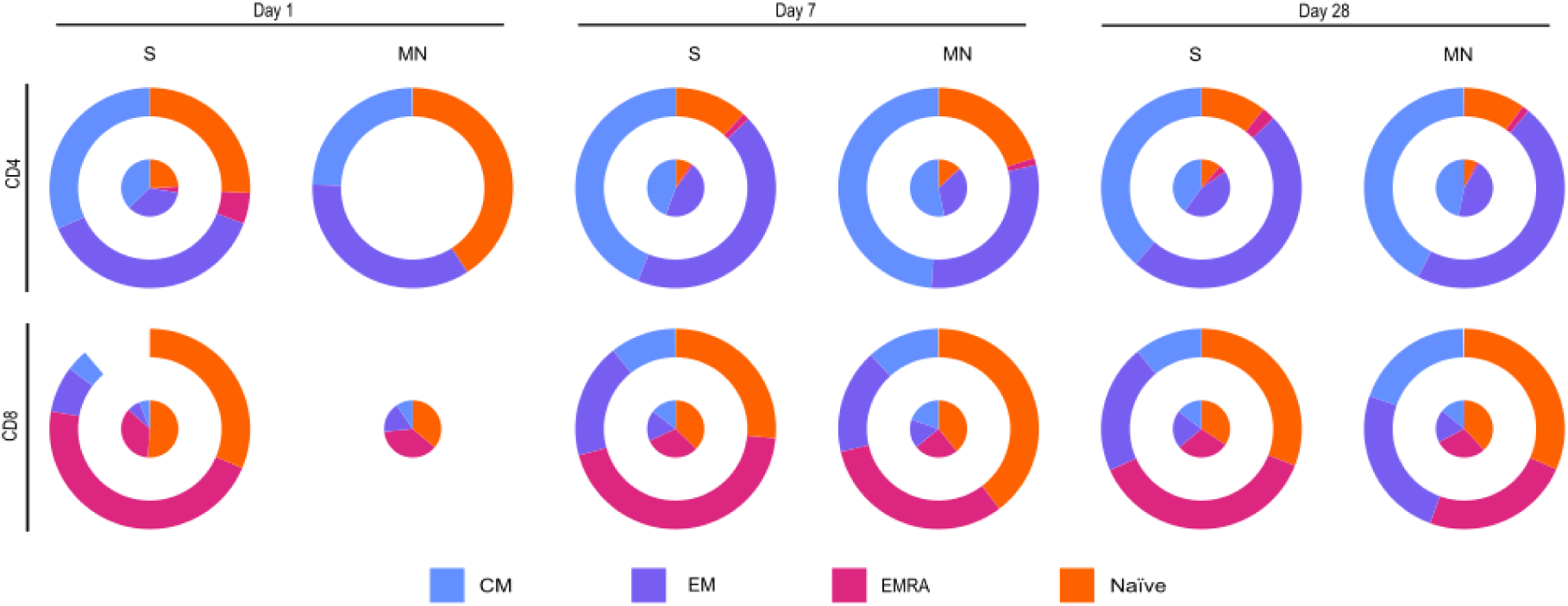
Mean distribution of memory subsets among SARS-CoV-2 specific CD4^+^ and CD8^+^ T cells. Memory subsets (CM: central memory; EM: effector memory; EMRA: effector memory re-expressing CD45RA) within stimulated AIM^+^ CD4^+^ T cells and AIM^+^ CD8^+^ T cells in 24 samples in three time points. Outer circle represents subsets for ivermectin-treated patients, and the inner circle for placebo-treated patients, top row depicts AIM^+^ CD4^+^ T cells and bottom row AIM^+^ CD8^+^ T cells. None of the comparisons of frequencies between treatment arms were significant as assessed by Wilcoxon-test.

The only significant difference observed between treatment arms was the frequency of S-specific naïve CD4^+^ T cells, which was lower in the IVM group at day 7 post-treatment (0.006% vs 0.036% of total CD4^+^ T cells adjusted p=0.02). No differences were found in the memory or the Th profile of the antigen-specific T cells between treatments (Figure 4).

### Ivermectin effect on main leukocyte subsets

We used an immunophenotyping panel to assess frequencies of different leukocyte populations in peripheral blood and their activation status (Supplementary Figure 5). A total of 99 cell subsets were analyzed. Of those, 21 cell subsets varied between time points within the placebo-treated group or between treatment groups by timepoint (unadjusted p-value <0.099, by T-test and Wilcoxon-test, respectively) and were selected for subsequent analyses. Cell subset frequencies, expressed as a percentage of all live cells, were analyzed using hierarchical clustering across all samples. While the clustering did not segregate the samples by treatment or time point (data not shown), cell subsets clustered according to common phenotypic features and changes overtime can be observed for certain cell clusters such as B cells and monocytes (Supplementary Figure 6).

Classical monocytes (CD14+CD16-), regulatory T cells (T-regs) and CM CD8^+^ T cells tended to increase from day 1 to day 7 after infection and remained higher at day 28 (Figure 6). Meanwhile, B cells and CD8^+^ Naïve T cells showed a more gradual increase from day 1 to day 28. Non-classical monocytes and the general monocyte-like population (gated based on scatter parameters and HLA-DR expression) increased from day 7 to day 28. In contrast, plasmablasts, CD56^+^CD16^-^ NK cells, EMRA CD8^+^ T and CD57^+^ CD8^+^ T cells (terminally differentiated and highly cytotoxic cells) showed a continued decrease from day 1 to day 28. Activated (HLADR+CD38+) CD4^+^, CD8^+^, and ɣδ T cells decreased from day 7 to day 28. Decreases in cell subsets likely reflect a contraction of cell subsets probably expanded during the acute infection and a return of activated cells to baseline levels.

**Figure 6.**
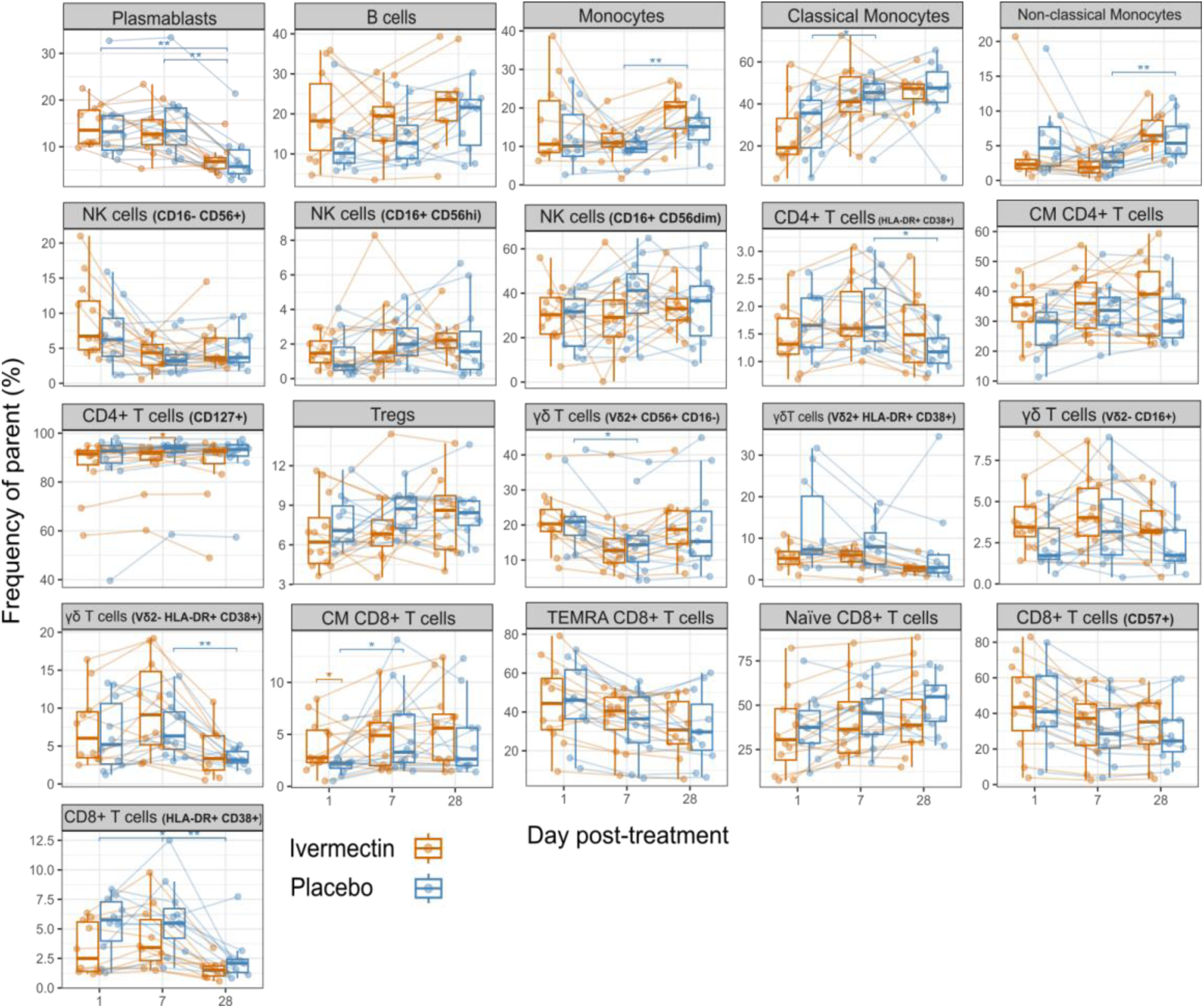
Kinetics of frequencies of leukocyte subsets. Frequencies of cell populations measured with a pan-leukocyte phenotyping panel in PBMC of the 24 study participants in three time points (data from same subject joined by lines). Cell subsets that exhibited any difference between time points within the placebo-treated group or between treatment groups at each time point (unadjusted p-value <0.099) are shown. For each time point box plots represent median and inter quartile range for 12 ivermectin-treated patients (orange) and 12 placebo-treated patients (blue), paired samples joined by lines. Statistically significant differences between time points among placebo-treated patients before adjusting appear in dark blue brackets and * indicate p <0.05, and ** p <0.01 from Wilcoxon-test (n=12). Statistically significant differences between treatment arms before adjusting appear in orange brackets and * indicates p <0.05 from Wilcoxon-test.

The only statistically significant differences before adjusting by multiple testing between treatment arms were a lower frequency of CD127^+^ CD4^+^ T cells (cells with high functional and survival capacity) in the IVM group at day 7 (87.57% of total CD4^+^ T cells vs 90.75%, unadjusted p=0.04) and a higher frequency of CM CD8^+^ T cells at day 1 (3.88% of total CD8^+^ T cells vs 2.32%, unadjusted p=0.04) (Figure 6). The latter reflecting heterogeneity in leukocyte frequencies between individuals at baseline.

### Ivermectin effect on coordination of cellular and antibody responses

A prompt humoral and cellular response has been related to control of the infection and milder COVID-19 [30]. We tested whether IVM-treated patients had a more coordinated response than placebo-treated patients using Spearman’s rho rank correlation coefficients for pairs of antigen-specific T-cell frequencies and antibody measurements. We did not find robust evidence of different correlation patterns in IVM and placebo-treated patients (Supplementary Figure 7).

### Ivermectin effect on whole blood transcriptomic profile

To have a more comprehensive systems-level insight, we analyzed the gene expression profile of blood cells at days 1, 4 and 7 post-treatment in both groups, independently. A total of 666 differentially expressed genes (DEG) were upregulated or downregulated at different study visits (day 4 and day 7) compared to baseline in the IVM-treated group, while 1,422 DEGs were found in the placebo group (Figure 7). A total of 23 Gene Ontology (GO) Biological Process pathways were upregulated or downregulated at different study visits (day 4 and day 7) compared to baseline in both treatment arms (Supplementary Fig 8). Downregulated pathways on day 7 compared to day 1 were related to viral and innate responses, whereas upregulated pathways on day 7 compared to day 1 were related to adaptive responses. A total of 58 GO Biological Process pathways were upregulated or downregulated at different study visits (day 4 and day 7) compared to day 1 in placebo group only and not in the IVM group, related to viral responses and cell cycle, respectively (Supplementary Fig 9).

**Figure 7.**
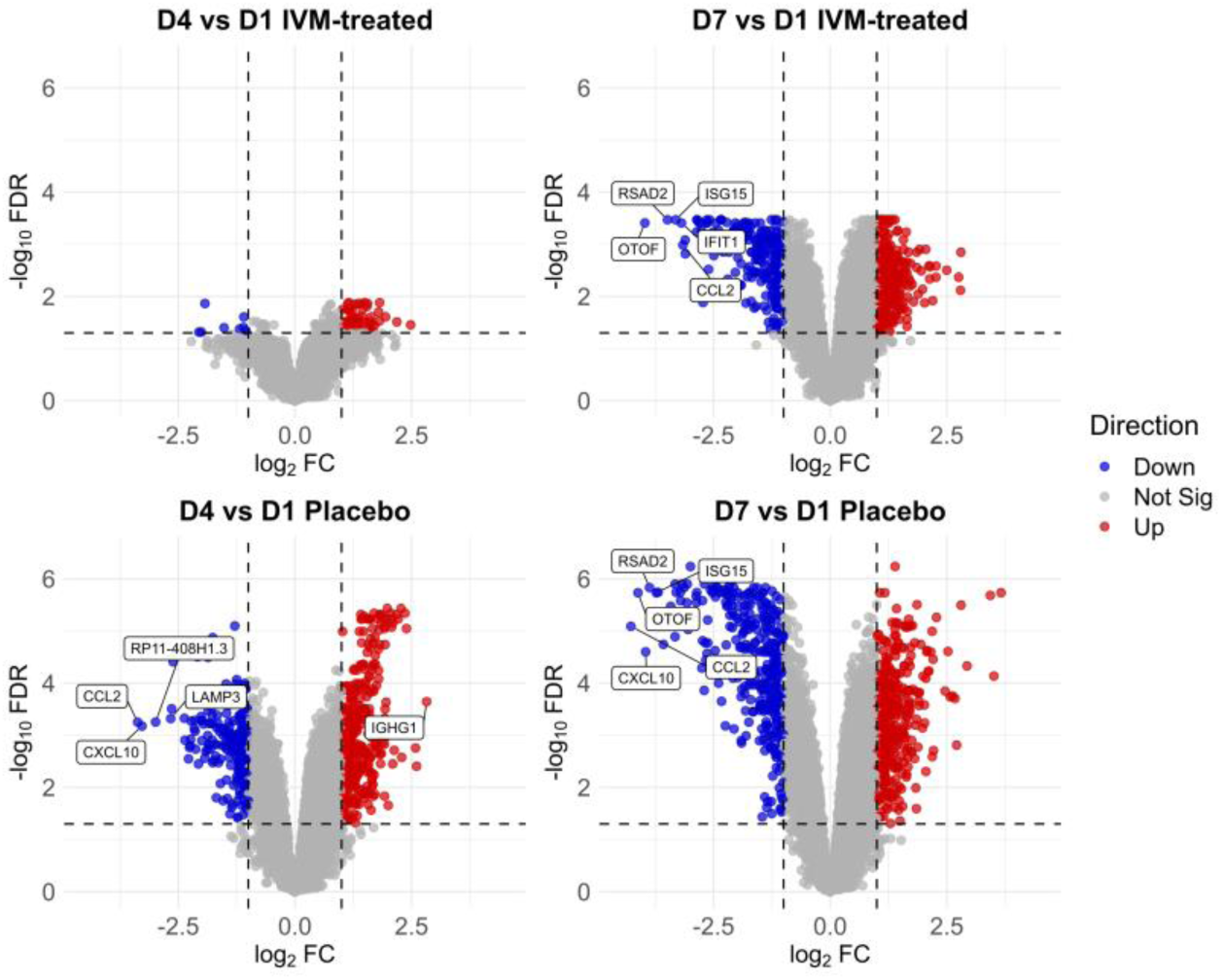
Volcano plots showing differentially expressed genes at days 4 and 7 compared to baseline in the ivermectin (IVM)-treated and placebo groups. Differential gene expression analysis was conducted using limma with voom transformation. No differentially expressed genes (DEG) were found at day 7 compared to day 4. Upregulated (red) and downregulated (blue) genes were defined as log2FC > 1 or < -1 with adjusted p < 0.05. Non-significant genes are in gray. Labels indicate the top 5 DEG with |log_2_FC| > 2.5.

**Figure 7.**
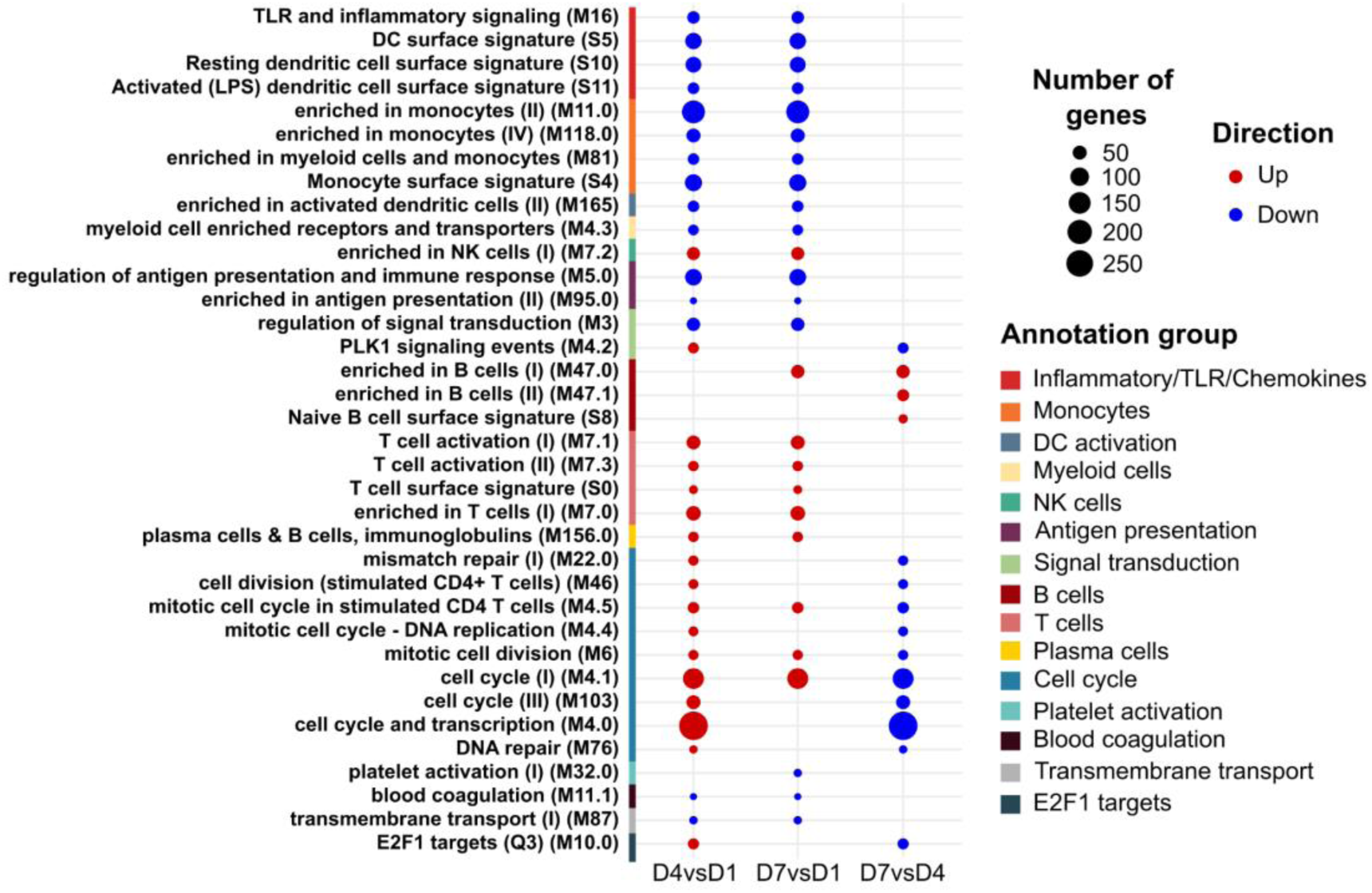
Commonly enriched BTMs in both treatment arms at different days post-treatment. Camera analysis and voom transformation were applied in both groups independently. BTMs with an adjusted p < 0.05 and number of genes > 20 were selected. A total of 36 enriched BTMs were found in common between treatment arms. Red dots indicate upregulated pathways and blue dots indicate downregulated pathways in the comparisons D4vsD1, D7vsD1, and D7vsD4. To be annotated (TBA) BTMs were removed since they lacked biological meaning.

A total of 36 blood transcriptional modules (BTMs) were upregulated or downregulated at different study visits (day 4 and day 7) compared to baseline (day 1) or at day 7 compared to day 4 in both treatment arms (Figure 8). Downregulated BTMs on day 4 and 7 compared to day 1 were related to innate immune responses and monocytes and dendritic cells, which seems contradictory with the increased classical monocyte frequencies after day 1 reported above. A BTM related to NK cells was upregulated at day 4 and day 7 compared to day 1 also in contrast to the immunophenotyping results. Upregulated BTMs were related to adaptive immune responses and cell subsets (T, B and plasma cells) as well as cell cycle BTMs, particularly at day 4. A total of 27 BTMs were upregulated or downregulated at different study visits in IVM-treated or placebo groups only (Supplementary Fig 10). In the IVM-treated group, pathways mainly related monocytes and DC were primarily downregulated on day 7 compared to day 4, whereas in the placebo group, pathways were primarily upregulated on day 7 (compared to day 1 and day 4) and were mainly related to adaptive responses (B and plasma cells).

**Figure 8.**
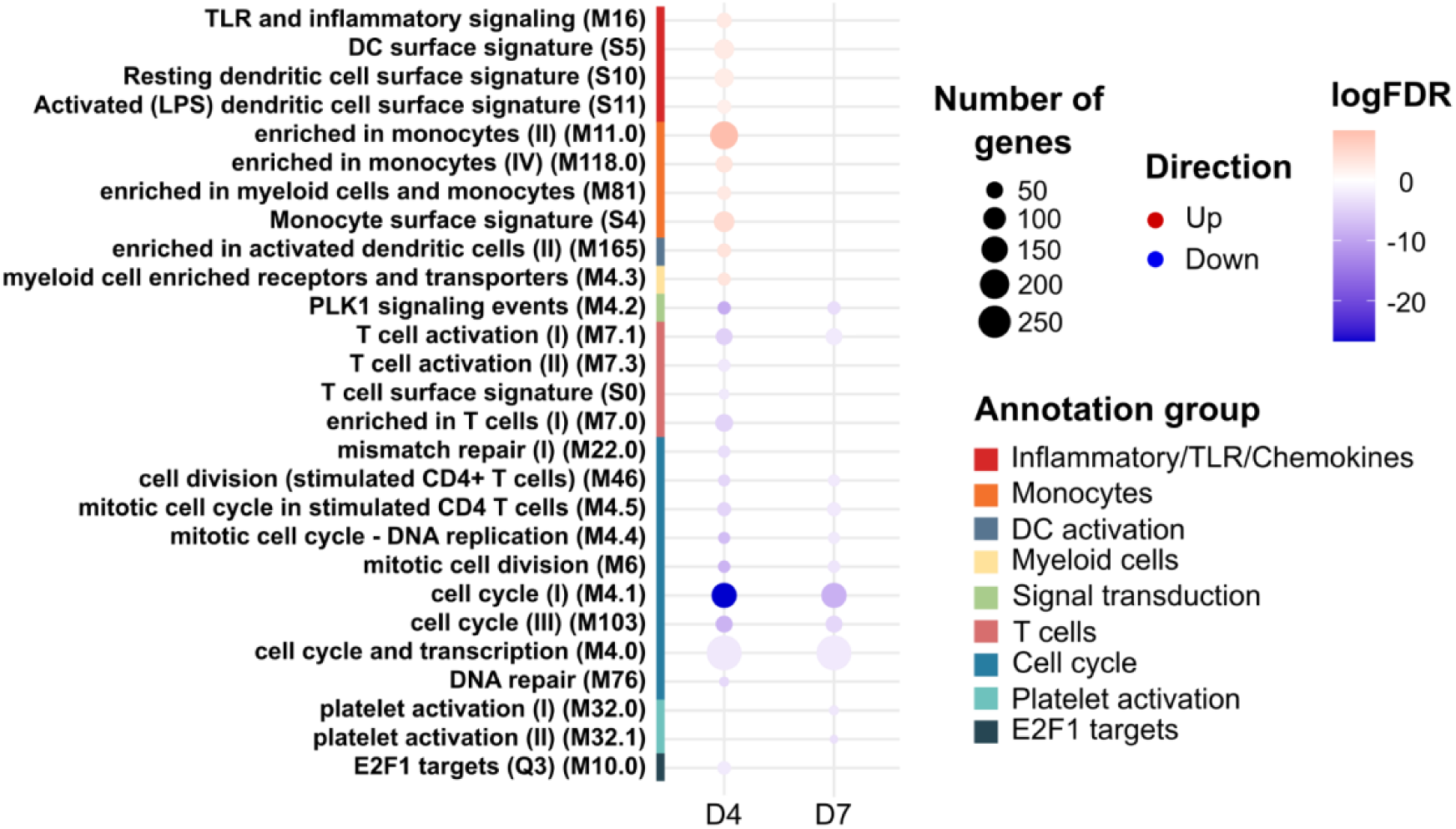
Enriched BTMs in the ivermectin (IVM)-treated group compared to placebo-treated groups at days 4 and 7 post-treatment. This analysis was done comparing ivermectin-treated (n=12) and placebo (n=12) groups on days 4 and 7 adjusting for the baseline. Camera method and voom transformation were applied. BTMs with an adjusted p < 0.05 and number of genes > 20 were selected. A total of 27 enriched BTMs were found. TBA BTMs were removed since they lacked biological meaning. Red dots indicate upregulated pathways while blue dots indicate downregulated pathways in the IVM-treated group compared to the placebo group

When comparing the IVM to the placebo arms, at day 4 and day 7, no DGE were detected (data not shown). However, we found 10 and 15 BTMs up- and downregulated, respectively, in the IVM-treated group (Figure 8). Downregulated BTMs were related to T cells and cell cycle, whereas upregulated BTMs were related to innate responses, mainly monocytes and DC, consistent with the above longitudinal transcriptional analysis. At day 7, fewer BTMs were enriched, and all 11 BTMs were downregulated in the IVM-treated group and were again related to T cells and cell cycle.

Overall, the transcriptomic profile suggests a strong innate and antiviral response at day 1 that decreases over the subsequent days, whereas a T and B cell response was being induced and probably peaking already at day 4. Although no DEGs were found when comparing treatment arms, differences in BTM and BTM uniquely enriched in the placebo group suggest less downregulation of innate and less induction of acquired responses in IVM-treated individuals.

## DISCUSSION

This prospective longitudinal study of humoral, cellular immune and transcriptomic components, conducted within the context of a randomized placebo-controlled clinical trial, did not reveal a significant immunomodulatory effect of IVM on responses to mild COVID-19. If there was an undetected effect, it must be minimal, in line with the results of clinical trials showing little or no pharmacological effect on COVID-19 outcomes.

Immune responses observed in COVID-19 participants were in line with what has been reported thus far. The main components of the adaptive immune response tended to increase within a few days in mild COVID-19, very early after infection [30]. Most COVID-19 patients seroconvert within 5-15 days post symptoms onset [31–35], developing IgM, IgA, and IgG to SARS-CoV-2 S and N proteins simultaneously. We saw that at day 4 (approximately 6 days after symptoms onset) levels of IgA and IgM started to increase and peaked at day 14, whereas IgG increased and peaked later. Early presence of antigen-specific antibodies correlates with milder disease, suggesting a role for adaptive immunity in infection control [30], and consistent with the decreased viral load by day 4 observed in the participants and the resolution of symptoms in a few days. Of note, we found a non-IgA responder, which had also been previously observed [30].

CD4^+^ and CD8^+^ T cells are detected in almost all SARS-CoV-2 infections [29], appearing within days and peaking at two weeks post-symptoms onset [32,36,37], with CD4^+^ T cells being more prominent than CD8^+^ T cells [38,39], which is consistent with our observations. Interestingly, in our study the Th1/Th2 balance was shifted towards a Th2 response, which is not in line with the literature, as CD4^+^ Th2 elevated responses have rarely been described [30,32,38,39]. Some studies have reported an underrepresentation of Th1 in severe cases [37], while a Th2-type response is usually a consequence of a dysregulated cytokine response [40], but in our study all cases were mild. Regarding the memory phenotype of SARS-CoV-2 specific T cells, CM and EM predominated in CD4^+^ T cells, and EM and EMRA in CD8^+^ T cells, similar to what has been described months after the infection [29]. Nevertheless, cellular responses were in general heterogeneous [30,32,38], which could be partially explained by the differences in hours post-symptoms onset at the first sample taken, which had a mean of 41.5 h with a 29.6 SD. Frequencies of other cell subsets changed over time, with an increase in classical monocytes by day 7 and non-classical monocytes by day 28, which could reflect return to baseline status after recruitment of monocytes to the airways compartments during the acute infection [41–43]. In contrast, plasmablasts decreased over time, probably reflecting the differentiation to antibody generating plasma cells and is aligned with the early antibody responses observed. Frequencies of activated CD4^+^, CD8^+^ and γδT cells decreased at day 28, suggesting a return to baseline activation levels after resolution of the infection. Cytokine kinetics also varied widely between patients, already decaying after day 1 post-treatment. All these changes are aligned with the viral load decrease by day 4, the quick resolution of symptoms, and the fact that no participant evolved to severe disease.

IVM did not exert a clear immunomodulatory effect on the SARS-CoV-2-specific antibodies studied, with only significantly higher levels of IgG against N at day 7 post-treatment. Pedroso et al. assessed the levels of antibodies against S and N proteins in blood and evaluated the in vitro production of antibodies by S-stimulated PBMCs in individuals taking different self-prescribed dosages of IVM [12]. They found that patients who had taken more than one IVM dose had lower antibody levels to S or N and their B cells produced less antibodies. However, patients who took multiple doses also reported higher symptomatology, and authors did not clarify whether IVM was taken before or after symptoms onset. It could be that patients with worse symptomatology took more IVM doses and the differences in antibodies reflected differences in severity of COVID-19 and the corresponding immune response rather than a dose-dependent effect of IVM. Besides, some of the patients had taken azithromycin as well, which has also been studied for immunomodulatory effects [44], and their results were not adjusted for multiple comparisons. We did not find here an effect for IVM on T cell responses either. Pedroso et al. also evaluated the in vitro production of Th1/Th2 cytokines and reported no differences across different IVM dosages[12]. Supporting a lack of effect of IVM on cytokine production, another study did not find significant differences in levels among six healthy volunteers who took a 0.25 mg IVM/kg single dose compared to six who took placebo [18]. This contrasts with previous studies in onchocerciasis and filariasis patients that found a potentiation of the cellular immune component [9], while a more recent study with ChPV-infected goats reported a decrease in CD8^+^ T cells [8].

Transcriptomic analyses did not allow us to detect any clear effects of IVM either. Observed gene expression responses reflect the kinetics of immune responses and the COVID-19 evolution of the participants. At days 4 and 7, in both treatment arms, there was a decrease in the innate and antiviral response as well as BTMs related to DC and monocytes compared to day 1, while there was an upregulation of BTMs related to adaptive immunity, including T and B cells and immunoglobulins as well as cell cycle, presumably from proliferating T cells. Nevertheless, an increased number of DEGs, GO pathways and BTMs related to the above immune responses were detected in the placebo group at day 4 and 7 compared to day 1. While no DEGs were found when directly comparing the two treatment arms, a few differences were detected using GSEA with BTMs, showing a stronger innate immune response on day 4 but decreased T cell responses in the IVM-treated group. We cannot discard that those differences are not due to a stronger response at day 1 in the placebo group resulting in increased changes to return to baseline status after resolution of infection. Our results are in line with the rest of immunological analysis performed to date, including Wilson *et al.* (2021), who observed several transcriptomic changes 4 h after IVM administration in human volunteers, effects that were no longer present by 24 h [18].

Our study has limitations, including the sample size that was modest, affecting statistical power and random distribution of the participants. Although no significant differences were found between groups, there could be imbalances at baseline in infection and subsequent immune responses and could have affected results between treatment groups. In addition, the administration of treatment was on average 44 h post-symptoms onset, and by day 4 the viruses were barely detected. Thus, the responses analyzed did not reflect acute infection but rather a phase of infection resolution and convalescence. Therefore, we cannot discard that IVM may alter immune responses to moderate or severe cases or if IVM is administered and evaluated earlier in the onset of infection. In addition, we have only assessed peripheral blood responses and not mucosal responses, which are relevant for this respiratory pathogen. However, the study has several strengths. It was performed in the context of a randomized clinical trial, the immune system characterization was done longitudinally and in great depth, covering innate and adaptive responses, pathogen specific and agnostic non-specific, and using complementary techniques. Additionally, a stringent exploratory analysis controlling for multiple comparisons was used.

While an antiviral activity of IVM has been already rejected at doses approved for human usage, a potential immunomodulatory activity that could improve the host response to infection, benefiting COVID-19 progression, has not been yet discarded. Our comprehensive longitudinal analysis of the immune responses and whole blood gene expression revealed no robust effects of IVM on SARS-CoV-2 early immune responses in mild COVID-19 cases. Overall, our results are congruent with the clinical trial results and evidences pointing towards a lack of effect of IVM on COVID-19. Nevertheless, we cannot discard that IVM has any immunomodulatory properties albeit if any, those are probably very modest and dependent on the experimental context, the pathogen targeted and related to the anti-pathogen effect.

## METHODS

### Ethics

The protocol of the clinical trial has been described elsewhere [25]. It was approved by the Spanish national ethics committee for drug research (Hospital Puerta de Hierro Majadahonda) and the Spanish Agency of Medicines and Medical Devices. All patients provided written informed consent and procedures were conducted in compliance with the latest version of the Helsinki Declaration and Good Clinical Practice.

### Study design, patient characteristics and samples obtained

The trial was conducted in Pamplona (Navarra, Spain). COVID-19 patients with a positive SARS-CoV-2 PCR and a maximum of 72 h of cough or fever, were invited to participate. The first patient was enrolled on July 31, 2020 and the last on September 11, 2020. Relevant exclusion criteria included having detectable IgG levels against SARS-CoV-2 measured by a rapid test, having had more than 72 h of fever or cough, COVID-19 pneumonia diagnosed by a physician or identified in a chest X-ray, age < 18 or > 60 years, and having any comorbidity that might pose a higher risk for progression to severe COVID-19.

At enrolment (day 1), patients received orally a single dose of 400 µg/kg of IVM or placebo along with an assessment of their vital signs. Laboratory parameters measured as common clinical practice were obtained for days 1, 7 and 14 post-treatment. Both nasopharyngeal and oropharyngeal samples were collected at days 1, 4, 7, 14, 21 and 28 post-treatment for PCR detection of SARS-CoV-2. Approximately 4 mL of blood samples in EDTA tubes were taken at days 1, 4, 7, 14, 21 and 28 post-treatment for plasma antibody and cytokine measurements, and 10 mL in heparin tubes at days 1, 7 and 28 for cellular assays. Finally, at days 1, 4 and 7 post-treatment, blood was collected in PAXGene tubes for RNA conservation and subsequent RNA sequencing.

Additionally, all patients were asked to fill in an online daily diary of symptoms from day 1 to 28 post-treatment. When there was noncompliance in answering the questionnaire on a given day, the previous day response was used.

### Quantification of antibodies against SARS-CoV-2

Plasma IgA, IgG and IgM levels (MFI) at days 1, 4, 7, 14, 21 and 28 post-treatment, against five SARS-CoV-2 antigens from the Wuhan strain were measured by quantitative suspension array technology (Luminex) in multiplex, as previously described [45]. Antigens included the full-length S (aa 1-1213 expressed in Expi293 and His tag-purified), its subregion S1 (aa 1-681, expressed in Expi293 and His tag-purified), both produced at the Center for Genomic Regulation, the subregion S2 (SinoBiological), RBD (StrepTag purified from the supernatant of lentiviral-transduced CHO-S cells cultured under a fed-batch system), and the N full-length protein (expressed in *E. coli* and His tag-purified). In addition, the N antigen of the HuCoVs 229E, HKU1, NL63 and OC43 (all expressed in *E. coli* and His tag-purified) were also included in the panel.

#### Coupling of proteins to microspheres

MagPlex® polystyrene 6·5 μm COOH-microspheres (Luminex Corp, Austin, TX, USA) were washed, sonicated and activated with Sulfo-NHS (N-hydroxysulfosuccinimide) and EDC (1-Ethyl-3-[3-dimethylaminopropyl] carbodiimide hydrochloride) (Thermo Fisher Scientific Inc., Waltham USA). Next, microspheres were washed and resuspended in 50 mM MES pH 5.0 (MilliporeSigma, St. Louis, USA). The recombinant proteins were then incubated with the microspheres at the optimal concentrations (from 10 to 50 μg/mL, depending on the antigen) and left at room temperature on a shaker for two hours. Coupled microspheres were resuspended in PBS with 1% BSA and 0.05% Tween 20 to covalently block the free carboxylic group (-COOH) absorbing most of the non-specific binding to secondary antibodies during assay steps [46] and heterophilic antibody binding seen in previous systems [47]. Microspheres recovery was quantified on a Guava® easyCyte™ Flow Cytometer (Luminex Corporation, Austin, USA). Equal amounts of each antigen-coupled microspheres were multiplexed and stored at 2000 microspheres/μL at 4°C, protected from light.

#### qSAT assay

Antigen-coupled microspheres were added to a 384-well Clear® flat bottom plate (Greiner Bio-One, Frickenhausen, Germany) in multiplex (2000 microspheres per analyte per well) in a volume of 90 μL of Luminex Buffer (1% BSA, 0.05% Tween 20, 0.05% sodium azide in PBS) using 384 channels Integra Viaflo semi-automatic device (96/384, 384 channel pipette). Two hyperimmune pools (one for IgG, and another one for IgA and IgM) were used as positive controls in each assay plate for QA/QC purposes and were prepared at 2-fold, 8 serial dilutions from 1:12.5. Pre-pandemic samples were used as negative controls. To quantify IgM and IgA responses, test samples and controls were pre-treated with anti-human IgG (Gullsorb) at 1:10 dilution, to avoid IgG interferences. The positive control, negative controls and test samples were prediluted 1:50 in 96 round-bottom well plates, and 10 μL were added to a 384-well plate using Assist Plus Integra device with 12 channels Voyager pipette. Final sample dilution was 1:500 for the three isotypes. Technical blanks consisting of Luminex Buffer (PBS with 1% BSA) and microspheres without sample were added in 4 wells to detect and adjust for non-specific microsphere signals. Plates were incubated for 1h at room temperature in agitation (Titramax 1000) at 900 rpm and protected from light. Then, the plates were washed three times with 200 μL/well of PBS-T (0.05% Tween 20 in PBS), using BioTek 405 TS (384-well format). Twenty-five μL of goat anti-human IgG phycoerythrin (PE) (GTIG-001, Moss Bio) diluted 1:400, goat anti-human IgA-PE (GTIA-001, Moss Bio) 1:200, or goat anti-human IgM-PE (GTIM-001, Moss Bio) 1:200 in Luminex buffer were added to each well and incubated for 30 min. Plates were washed and microspheres resuspended with 80 μL of Luminex Buffer. Before acquisition on the Flexmap 3D® reader, plates were sonicated 20 seconds. At least 50 microspheres per analyte per well were acquired, and MFI was reported for each isotype-antigen pair.

### Quantification of cytokines/chemokines and growth factors

The Cytokine Human Magnetic 30-Plex Panel from Invitrogen™ was used to measure the concentrations of 30 analytes in serum from patients at days 1, 4, 7, 14, 21 and 28 post-treatment: epidermal growth factor (EGF), fibroblast growth factor (FGF), granulocyte colony-stimulating factor (G-CSF), granulocyte-macrophage colony-stimulating factor (GM-CSF), hepatocyte growth factor (HGF), VEGF, tumor necrosis factor (TNF), IFN-α, IFN-γ, IL-1RA, IL-1β, IL-2, IL-2R, IL-4, IL-5, IL-6, IL-7, IL-8, IL-10, IL-12(p40/p70), IL-13, IL-15, IL-17, IP-10, MCP-1, MIG, macrophage inflammatory protein (MIP)-1α, MIP-1β, regulated on activation normal T cell expressed and secreted (RANTES) and eotaxin. Samples were tested in single replicates following a modification of the manufacturer’s protocol explained elsewhere[48]. Each plate included 16 serial dilutions (2-fold) of a standard sample provided by the vendor and, two blank controls. Samples were acquired on a Luminex® 100/200. The concentration of each analyte was obtained by interpolating the MFI to a 5-parameter logistic regression curve and reported as pg/mL using the drLumi R package (6). Limits of quantification (LOQ) were estimated based on cutoff values of the 30% coefficient of variation (CV) of the standard curve for each analyte. Values below the lower LOQ (LLOQ) were assigned a random value between the LLOQ and the LLOQ/2. Values above the upper LOQ (ULOQ) were assigned a random value between the ULOQ and the ULOQx2.

### Cellular assays

PBMCs were isolated from venous blood samples taken at day 1, 7 and 28 after treatment by density-gradient centrifugation using Ficoll-Paque and cryopreserved in heat-inactivated fetal bovine serum (HI-FBS) with 10% dimethyl sulfoxide (DMSO) and stored in liquid nitrogen until shipping. Frozen PBMCs were then thawed in 5 ml of TexMACS medium (Miltenyi) with 1% penicillin/streptomycin (P/S) (ThermoFisher) and 12.5 U/ml DNase (Benzonase® Nuclease, Merck). Cells were washed with TexMACS medium + 1% P/S and counted on the Guava® easyCyte™ Cytometer (Luminex) at a 1:40 dilution with Guava ViaCount™ Reagent (Cytek Biosciences). For immunophenotyping, 0.5-1×10^6 cells were stained with the antibody panel described in Supplementary Table 1 and fixed with a 1% paraformaldehyde (PFA) solution. The rest of PBMC were resuspended at 2×10^6 cells/mL with culture medium in 50 ml tubes and rested overnight (ON) at 37°C and 5% CO_2_. After ON resting, cells were counted again and resuspended at 5×10^6^ cells/mL and 200 µL per condition were plated in a 96 well round bottom plates (VWR) for the AIM assay to assess SARS-CoV-2-specific T cells. Rested PBMCs were stimulated with peptide pools comprising the protein sequences of the S (PepTivator® SARS-CoV-2 Prot_S Complete, Miltenyi), N (PepTivator® SARS-CoV-2 Prot_N, Miltenyi) and M (PepTivator® SARS-CoV-2 Prot_M, Miltenyi). Peptides were 15 amino acids long with an 11-amino acid overlap, dissolved in sterile water and used at a 1 µg/mL/peptide concentration. The N and M peptide pools were combined into a single stimulus. The mitogen phytohemagglutinin (PHA) (Merck) was used as a positive control at a 10 µg/ml concentration. Stimulations were prioritized in the following order: negative control (only media background control), S, MN and PHA. Plates were incubated undisturbed at 37°C, 5% CO_2_ for 24 h followed by staining with the antibody cocktail described in Supplementary Table 1 & 2.

The staining protocol was the same for the immunophenotyping and AIM antibody panels. Cells were stained with a 1:1000 dilution of the viability dye (eBioscience™Fixable Viability Dye eFluor™ 506, ThermoFisher) for 30 min at 4°C to exclude dead cells from the analysis. Afterwards cells were washed twice with PBS and stained with the antibody cocktail for 30 min at 4°C. All staining was done at a final volume of 50 µL per well. After staining, cells were washed twice with PBS + 2% FBS, fixed overnight with a 1% PFA solution and acquired in a BD LSRFortessa™ five-laser flow cytometer. Cytometer performance was tracked through acquisition of Sphero™ Rainbow Calibration Particles (8 peaks) (BD Biosciences). Data analysis was conducted in FlowJo™ v10.8 (BD Life Sciences), population gates were established through fluorescence minus one (FMO) controls. Gating strategies are shown in Supplementary Figures 3 & 5. Antigen-specific T cells were defined as the background-subtracted frequencies of AIM^+^ CD4^+^ (CD137^+^ OX40^+^) and AIM^+^ CD8^+^ (CD137^+^ CD69^+^) in each stimulation condition.

### Transcriptomic analyses

#### Extraction of RNA

RNA extraction from whole blood samples was performed using the Qiagen Blood miRNA Purification Kit, following the manufacturer’s protocol. Whole blood was centrifuged to pellet the cells, washed with water, digested with proteinase K and homogenized via the provided PAXgene Shredder spin columns. Isopropanol was added to optimize RNA binding conditions, and RNA was bound to the silica membrane in PAXgene RNA spin columns. After DNase digestion to remove genomic DNA, the RNA was washed and eluted with the provided buffer.

#### RNA-sequencing

Library preparation and sequencing were performed at Centre de Regulació Genòmica (CRG, Spain). Universal Plus mRNA-Seq with NuQuant, Human Globin AnyDeplete, UDI-C kit for Illumina was used to process the samples. RNA sequencing was performed with the Illumina NextSeq 2000 using single-end 50 bp reads and achieving an average sequencing depth of 40 million reads per sample.

#### RNA-Seq data processing

FASTQ pre-processing (including quality control and quality trimming), RNA-Seq alignment and read quantification was performed. FastQC v.0.1.2.1 was used for raw read quality control. Low-quality read trimming was conducted using sickle v.1.33. RNA-Seq reads were aligned against human reference genome GRCh38 using RNA-Seq aligner STAR v.2.7.10b with settings ‘-outSAMunmapped Within’ and ‘-ouSAMtype BAM SortedByCoordinate’. GENCODE human genome annotation v.43 was used for raw read counting. The gene read quantification was performed using htseq-count v.2.0.3 with settings ‘-r pos’ and ‘-t gene’.

#### Differential gene expression (DGE) analyses

For downstream analysis of the transcriptomics data, 20,722 genes that had a minimum of 10 read counts in at least 12 samples were included, excluding globin-related genes. The trimmed means of M values (TMM) method was used to produce normalization factors correcting raw counts for different library sizes.

DGE analysis was performed using the limma-voom workflow as implemented in the limma v.3.56.2 package. Two DGE analyses were conducted: 1) assessment of changes within treatment arms from day 1 to day 7, with day as the predictor variable and contrast matrices for day 4 vs day 1, day 7 vs day 4 and day 7 vs day 1, and 2) comparison between treatment arms at days 4 and 7, with treatment as the predictor variable and subtracting the baseline to adjust for baseline individual variability.

For both analyses, a linear model without an intercept was performed, with RNA availability as a covariate since quantity of RNA was associated with duplicates detected in the FastQC. The lmFit function was used to fit the model for each gene, and contrasts were estimated with contrasts.fit and eBayes functions. P-values were adjusted for multiple comparisons using the Benjamini and Hochberg (BH) method to control the False Discovery Rate (FDR). Adjusted P-values < 0.05 and log2 fold-change (log2FC) > 1 were used to select DEGs.

#### Functional gene expression analyses

Gene set analyses using BTMs and voom transformation and camera, both from the limma package, were conducted to assess gene set differences directly, without first conducting a DGE analysis. To Be Annotated (TBA) BTMs were removed since they lacked biological meaning. In addition, clusterProfiler package v.4.8.2 and the GO database were used for enrichment analysis. Gene symbols were converted to ENTREZ ID using the org.Hs.eg.db v.3.17.0 package. Enrichment analysis of DGEs was then performed using the enrichGO () function with settings ‘ont=” BP”’. In both analyses, an FDR cutoff of BH adjusted P-values < 0.05 and number of genes > 20 were used to select significant gene sets.

### Statistical analysis

Differences between treatment arms for baseline comparisons were tested using T-test for normally-distributed numeric variables and Wilcoxon test for non-normally distributed ones-tested by Shapiro-Wilk test-; and a Chi-square test of independence for categorical variables.

Measurements of antibody levels (MFI) and cytokine concentrations (pg/mL) were log_10_-transformed, given that antibody levels and cytokine concentrations are well approximated by a log-normal distribution. To explore the antibody and cytokine immune response to SARS-CoV-2 independently of treatment we used the placebo-treated samples and compared levels between time points using Linear Mixed Models (LMM) including a random intercept to account for repeated measurements with participant ID as the grouping factor. Differences in geometric mean levels of antibodies between timepoints relative to baseline were estimated using coefficients for fixed effect indicator regressors encoding time point. P-values were obtained from t-tests on the coefficients corrected by Satterthwaite’s method for degrees of freedom and BH for multiple comparisons.

To compare levels of antibodies and cytokines between treatment arms across multiple time points we performed a likelihood ratio test (LRT) comparing full and reduced models using niched LMMs. This allowed us to reduce the number of tests performed as opposed to doing paired tests for each time point. Reduced models only contained fixed effect time point indicator regressors. By contrast, full models included an interaction between the time point indicator regressor and the treatment, therefore the model estimated a distinct mean per time point and per treatment arm. Day 1 was excluded as treatment had just been given and no effect was expected. The LRT indicated whether the full model provided a better fit for the data, which was then interpreted as influencing significantly the levels of that particular analyte. A post-hoc analysis using Wilcoxon test was then performed to see which particular time point showed significantly different levels between arms.

To analyze the cellular immune response, we used a different approach, as data obtained from flow cytometry is presented as frequencies, which do not usually follow a normal distribution. Therefore, a paired Wilcoxon test was used to compare levels in the placebo group between the three time points, and between treatment arms for each time point.

95% CIs were provided. FDR was accounted for by adjusting p-values with BH method within each set of identical hypotheses tested. An adjusted p-value of ≤0.05 was considered statistically significant.

When we compared results from various assays using different units within the same plot, we calculated a fraction peak. Here, the peak represented the maximum value of the given measurement across all time points, ensuring a standardized range of zero to one regardless of the original units.

We performed the statistical analysis in R version 4.2.1 (packages dplyr, tidyr, ggplot2, lmerTest, lme4, vtable and corrplot).

## DATA AVAILABILITY STATEMENT

Sequencing data have been deposited in GEO under accession code GSE290188. Immune phenotyping and antibody data used for the analysis are archived on Dora repository (doi xxxx). The analysis code can be found at Dora repository (doi xxxx).

## ACKNOWLEDGEMENTS

We thank the patients for their participation in the study and the clinical and lab team in Navarra and the lab team in ISGlobal. We are very grateful to Juan C. Gabaldon-Figueira, the nursing staff of the emergency room, the technicians of the microbiology lab of the Clínica Universidad de Navarra as well as people from the Biobank for their dedication to SAINT study, and to Laura Puyol for logistics coordination. We are grateful also to Natalia Rodrigo Melero (CRG), Daniel Parras and Pau Serra (IDIBAPS), for their contribution on protein production, and to the Cytometry and cell sorting core facility of the Institut d’Investigacions Biomèdiques August Pi i Sunyer (IDIBAPS) for the technical help.

## SUPPORTING INFORMATION CAPTIONS

### Supplementary Figures

**Supplementary Figure 1. Kinetics of antibody levels to N antigen from human coronaviruses causing the common cold since day of treatment.** Log-10 transformed levels (median fluorescence intensity, MFI) of IgA, IgG, and IgM against the N antigen of HKU1, NL63, OC43 and 229E coronaviruses measured in 24 patients across six time points (data from same individual are joined by lines). For each timepoint boxplots represent median and interquartile range MFI for ivermectin-treated patients (orange) and placebo-treated patients (blue).

**Supplementary Figure 2**. **Kinetics of cytokine concentrations since day of treatment.** Levels (pg/mL) of cytokines measured in 24 patients across six time points (data from same individual are joined by lines). For each timepoint boxplots represent median and interquartile range pg/mL for ivermectin-treated patients (orange) and placebo-treated patients (blue). Represented are the 16 out of the 30 cytokines analyzed that did not show statistical differences between timepoints.

**Supplementary Figure 3. Gating strategy for the AIM assay.** Example of the flow cytometry staining and gating strategy for the AIM panel. Gates were previously defined using fluorescence minus one (FMO) controls. **A.** The time vs FSC-A gate was used to exclude any acquisition irregularities. FSC-H and FSC-A were used to discriminate singlets and SSC-A vs FSC-A to select lymphocytes. Monocytes, B cells and dead cells were excluded by selecting CD14^-^, CD19^-^ and eFluor506-cells. From live cells, CD3+ cells were identified, with further gating of CD4^+^ and CD8^+^ T cells. Within these populations, memory subsets were identified through CCR7 and CD45RA expression: central memory (CM, CD45RA^-^ CCR7^+^), effector memory (EM, CD45RA^-^ CCR7^-^), effector memory re-expressing CD45RA (EMRA, CD45RA^+^ CCR7^-^) and naïve cells (CD45RA^+^ CCR7^+^). Additionally, T-helper phenotype was evaluated in CD4^+^ and CD8^+^ T cells, through CD183 (Th1) and CD194 (Th2) expression. Activated CD4+ and CD8+ T cells were defined by the expression of activation-induced markers (AIM) CD137^+^ OX40^+^ and CD137^+^ CD69^+^, respectively. Within AIM^+^ CD4+ and AIM^+^ CD8+ T cells, memory and T-helper phenotypes were identified through the same previously described gates. **B.** Representative dot plots showing the AIM expression in CD4^+^ (CD137^+^ OX40^+^) and CD8^+^ (CD137^+^ CD69^+^) T cells for each condition: unstimulated (US), spike (S), membrane (M) + nucleocapsid (N) and positive control (PC).

**Supplementary Figure 4. Kinetics of SARS-CoV-2 MN-specific T cells since day of treatment.** Frequencies of AIM+ CD4^+^ and CD8^+^ T cell subsets after MN stimulation and background subtraction. **A**. Frequencies of MN-specific CD4^+^ and CD8^+^ T cell populations. **B**. Frequencies of MN-specific central memory (CM), effector memory (EM), effector memory re-expressing CD45RA (EMRA) CD4^+^ and CD8^+^ T cell populations. **C.** Frequencies of Th1 and Th2 cells within MN-specific CD4^+^ and CD8^+^ T cells. For each timepoint box plots represent median and inter quartile range % for ivermectin-treated patients (orange) and placebo-treated patients (blue), paired samples are joined by lines, except for Day 1 for which it was not possible to test samples from the ivermectin-treated participants due to limited cell availability. For day 1: 4 placebo samples and none IVM were included; for day 7: 9 samples for each treatment arm were included; and for day 28: 11 samples for each treatment arm were tested. Statistically significant differences between time points among placebo-treated patients before adjusting appear in dark blue brackets and * indicate p < 0.05 from Wilcoxon-test (n=12). None remained significant after adjustment with BH (n=12). No statistical differences between treatments for each time point were found.

**Supplementary Figure 5. Gating strategy for the immunophenotyping assay.** Example of the flow cytometry staining and gating strategy for the immunophenotyping panel. Gates were previously defined using fluorescence minus one (FMO) controls. **A.** The time vs SSC-A gate was used to exclude any acquisition irregularities. FSC-H and FSC-A were used to discriminate singlets, and dead cells were excluded by an amine reactive dye. Monocytes were identified as HLA-DR^+^ and subsequently gated into three main subsets through CD14 and CD16 staining (classical CD14^+^ CD16^-^, intermediate CD14^+^CD16^+^ and non-classical CD16^+^CD14^dim^). **B.** Lymphocytes were selected from a boolean gate of monocyte-live cells, through SSC-A and FSC-A. NKT-like cells are defined through CD3 and CD56 expression. Subsequent gating discriminates CD3^+^ cells, followed by identification of Vδ2^+^ and Vδ2^−^ γδ T cells, CD4^+^ and CD8^+^ T cells. Within CD4^+^ T cells, Tregs are defined by being CD127^-^ and CD25^hi^. Within CD3^-^ cells, five NK cell subsets are defined by CD56 and CD16 expression (CD56^hi^CD16^−^, CD56^dim^CD16^−^, CD56^hi^CD16^+^, CD56^dim^CD16^+^ and CD56^−^CD16^+^) and B cells are identified as CD19^+^ HLA-DR^+^. Within B cells, plasmablasts are discriminated by a high CD38 expression. **C.** After negative gating on all lineage markers (CD3-CD19^-^CD14^-^CD16^-^ CD56^-^), dendritic cells (DCs) are defined by HLA-DR^hi^ expression. **D, E.** The expression of CCR7, CD45RA, CD57 and CD127 is examined within CD4^+^, CD8^+^, Vδ2^+^ and Vδ2^−^ γδ T-cell subsets to define memory and differentiation; HLA-DR and CD38 are evaluated in the same subsets to assess activation. Lastly, the expression of CD16, and CD16 in conjunction with CD56 and CD57 was evaluated within Vδ2^+^ and Vδ2^−^ γδ T-cells.

**Supplementary Figure 6. Heatmap of leukocyte subset frequencies.** Cell subset frequencies (calculated as a percentage of all live cells) were transformed to Z-scores and are shown as rows, colored by categories based on common features. Each column represents an individual sample, ordered by time point (Day) and treatment group. Rows were clustered using hierarchical clustering with Euclidean distance and the Ward.D method, and the color scale represents Z-scores ranging from -2 (low, white) to 2 (high, dark blue) z-score values. Only cell subsets that exhibited any difference between time points within the placebo-treated group or between treatment groups at each time point (unadjusted p-value <0.099, Wilcoxon-test) are shown (p< 0.099, statistical test).

**Supplementary Figure 7**. **Correlations of S-specific CD4^+^ T cell frequencies and RBD IgG levels.** Correlations of S-specific CD4^+^ T cells (frequencies of AIM^+^ T cells after spike stimulation and subtraction of background control) and RBD IgG levels (log10MFI) among 12 ivermectin-treated patients (orange) and 12 placebo-treated patients (blue). Rho spearman correlation values and their p-values are provided according to treatment.

**Supplementary Figure 8**. **Enriched GO pathways at day 4 and day 7 compared to baseline in common between treatment arms.** ClusterProfiler package and the GO database were applied in both groups, independently. GO pathways with an adjusted p < 0.05 and number of genes > 20 were selected. A total of 23 enriched GO pathways were found different between treatment arms.

**Supplementary Figure 9**. **GO pathways differently enriched between treatment arms.** CusterProfiler package and the GO database were applied in both groups, independently. GO pathways with an adjusted p < 0.05 and number of genes > 20 were selected. A total of 59 enriched GO pathways were found different between treatment arms.

**Supplementary Figure 10**. **BTMs differently enriched between treatment arms.** For this approach, camera method and voom transformation were applied in both groups, independently. BTMs with an adjusted p < 0.05 and number of genes > 20 were selected. A total of 27 enriched BTMs were found different between the treatment arms. TBA BTMs were removed since they lacked biological meaning.

## Supplementary Tables

**Supplementary Table 1.** Antibody panel for the Immunophenotyping assay

**Supplementary Table 2**. Antibody panel for the Activation-Induced Markers (AIM) assay

